# Exosomal miR-23a release coincides with astrocytic Calcineurin activation, protects neurons from apoptosis by regulating NOXA in Parkinson’s disease models and is enriched in the plasma of patients with Parkinson’s disease

**DOI:** 10.64898/2026.09.09.750353

**Authors:** Pallabi Bhattacharyya, Kusumika Gharami, Atanu Biswas, Subhas C. Biswas

## Abstract

Parkinson’s disease (PD) is characterized by a progressive loss of dopaminergic (DA) neurons in the substantia nigra pars compacta and degeneration of their projections in the striatum (STR) of the brain. Astrocytes are recognized as active contributors to PD pathogenesis, yet the role of calcineurin (CN), a Calcium/calmodulin-dependent phosphatase which is well characterized in neurons, remains unexplored in astrocytes in the context of PD. Here, we show that CN protein expression is upregulated in the STR of MPTP-treated mice, coinciding with DA neurodegeneration and astrogliosis. In human astrocytoma cells and primary rat astrocytes, the neurotoxin Rotenone induced rapid intracellular Calcium elevation and CN activation. Unexpectedly, this was accompanied by a decrease in intracellular levels of miR-23a, an astrocyte-enriched microRNA previously linked to CN signalling in cardiomyocytes. We found that this decrease reflected active exosomal export: Rotenone increased both exosomal miR-23a levels and total exosome number, while blocking exosome biogenesis with GW4869 restored intracellular miR-23a levels. Functionally, both astrocyte-derived exosomes and miR-23a overexpression protected neuronal cells (SH-SY5Y, PC12) against Rotenone-, 6-OHDA-, and MPP+-induced toxicity. Mechanistically, miR-23a showed direct binding to the 3′UTR of the pro-apoptotic BH3-only protein NOXA, thereby suppressing its expression, subsequently attenuating caspase-3 activation. Finally, exosomal miR-23a levels were significantly elevated in plasma from PD patients compared with age-matched controls, supporting clinical relevance. Together, these findings identify a previously unrecognized astrocyte-to-neuron communication axis during PD pathogenesis, in which astrocytic CN activation accompanied by exosomal miR-23a release represents a neuroprotective stress-responsive pathway with potential biomarker and therapeutic relevance in PD.

## 1. Introduction

Neurodegenerative disorders (NDDs) constitute a significant global health burden, with Parkinson’s disease (PD) ranking as the second most prevalent disease in this group. Approximately 1–4% of individuals older than 60 years are estimated to be affected by PD, which is characterized by one or all of the four cardinal signs of motor disorder-bradykinesia, resting tremor, muscular rigidity, and postural instability (Kalia and Lang 2015; Rocha et al. 2018). The neuropathological hallmarks of PD include progressive depletion of dopaminergic (DA) neurons in the substantia nigra pars compacta (SNpc), together with the formation of Lewy bodies, which are intracellular deposits containing aggregated α-synuclein, a protein strongly implicated in PD-associated neurodegeneration (Lees et al. 2009). Although majority of research on NDDs have primarily concentrated on neurons, reports over the last two decades is making it increasingly clear that NDDs are not just neuropathies; non-neuronal glial cells, especially astrocytes, are also affected or involved in the pathogenesis and progression of NDDs (Saha et al. 2020; Sarkar et al. 2025; Uemura et al. 2026).

Astrocytes maintain CNS structural and homeostatic integrity, supporting blood–brain barrier function, neurotransmitter recycling, and metabolic support to neurons (Abbott 2002; Ransom and Ransom 2012; Sofroniew and Vinters 2010; Vasile et al. 2017). In NDDs, astrocytes undergo “reactive gliosis” or “astrogliosis,” characterized by somatic- and process hypertrophy alongside increased expression of intermediate filament proteins like GFAP, vimentin, inflammatory cytokines, and oxidative stress markers (Eddleston and Mucke 1993; Eng et al. 2000; Ridet et al. 1997). This reactive phenotype is frequently associated with elevated intracellular calcium (LaFerla 2002; Takuma et al. 2004; Zawadzka and Kaminska 2005), which often activates calcineurin (CN), a Ca²⁺/calmodulin-dependent serine/threonine phosphatase highly expressed in neural tissue (Klee et al. 1979). It is composed of a catalytic CN-A subunit (∼59–61 kDa) and regulatory CN-B subunit (∼19 kDa) (Crabtree and Olson 2002; Feske et al. 2003). While the full length CN-A is ∼ 59-61 kDa, hyperactivation of CN can lead to proteolytic removal of a critical autoinhibitory domain located near the C-terminus, resulting in a truncated CN fragment in the ∼ 45-48 kDa size range (Pleiss et al. 2016; Kraner et al. 2024). Although (Norris et al. 2005) established a role for CN signalling in astrocyte reactivation, its downstream targets in astrocytes remain poorly characterized. To our knowledge, astrocytic Ca²⁺/CN signalling and its cellular consequences has not previously been explored in the context of PD.

Exosomes (30–150 nm) are a class of extracellular vesicles (EVs) that facilitate communication between astrocytes and neurons (Datta Chaudhuri et al. 2020). They modulate recipient-cell function through surface receptor interactions or by transferring their cargo via endocytosis or membrane fusion (Krämer-Albers and Werner 2023). Exosomal cargo includes diverse biomolecules, notably non-coding RNAs such as microRNAs (miRNAs) (Bhattacharyya et al. 2026; Nolte-’t Hoen et al. 2012; Valadi et al. 2007). miRNAs are small (20–22 nt) regulatory RNAs that, via Argonaute-containing complexes, bind complementary 3′UTR sequences to suppress translation or promote degradation of target mRNAs (Bartel 2018; Iwakawa and Tomari 2022; Filipowicz et al. 2008). Dysregulated miRNA expression is increasingly linked to disease pathogenesis given each miRNA’s capacity to regulate hundreds of target genes (Kim et al. 2025). The selective enrichment of miRNAs in exosomes has been shown to depend on specific sequence or post-translational modifications, suggesting that the loading of nucleic acids into exosomes is a regulated process (H. Lee et al. 2019; Villarroya-Beltri et al. 2013).

In NDDs, several brain-enriched miRNAs have been implicated in disease pathogenesis. In our previous study, we demonstrated that exosome-associated miR-128, a brain-enriched miRNA, was significantly reduced in the plasma of patients with PD and in cellular models of PD, while restoration of miR-128 protected neurons from 6-OHDA-induced apoptosis and preserved synaptic integrity (Bhattacharyya et al. 2023). On the other hand, miR-23a is reported to be astrocyte-enriched (Smirnova et al. 2005) and has been proposed as a candidate astrocyte marker (Gioia et al. 2014). Recently, (Barbagallo et al. 2020) found that miR-23a is upregulated in PD patient serum. Mechanistically, miR-23a upregulation is CN/NFATc3-dependent in cardiac hypertrophy (Lin et al. 2009), and (Hudson et al. 2014) reported in skeletal muscle atrophy, CN-dependent miR-23a is selectively exported via exosomes.

In this study, we demonstrate for the first time that CN expression is upregulated in the STR of MPTP-mouse models of PD which is accompanied by increased astrogliosis and DA neurodegeneration. Following Rotenone exposure, activation of the Ca²⁺/CN occurs in astrocytes accompanied by the exosomal release of astrocyte-enriched miR-23a. Astrocyte-derived exosomal miR-23a subsequently protects neurons from BH3-only protein (NOXA, PUMA) associated apoptosis under neurotoxic stress. These findings identify a previously unrecognized mechanism of astrocyte–neuron communication that contribute to neuronal protection, potentially during the early stages of PD-associated neurotoxic stress.

## 2. Materials and Methods

### 2.1. Acute MPTP mouse model

Adult male C57BL/6 mice (n = 4 per group) were divided into two experimental groups: a WT (control) group receiving saline injections, and an MPTP-treated group. MPTP was given intraperitoneally at 30 mg/kg of body weight per injection, administered twice, 16 h apart. Animals from both groups were euthanized on day 8 post-MPTP administration, following established protocols (Haobam et al. 2005; Mondal et al. 2023; Muralikrishnan and Mohanakumar 1998).

### 2.2. Immunohistochemistry and confocal imaging

Mice were deeply anesthetized with sodium pentobarbital and perfused transcardially with PBS, followed by 4% paraformaldehyde (PFA) prepared in PBS (pH 7.4). Brains were subsequently removed, post-fixed in 4% PFA for 24 h, and cryoprotected in 30% sucrose in PBS for 48 h. Coronal sections (40 μm thick) were prepared using a cryostat (SLEE MEV, GmbH) and maintained as free-floating sections in PBS until immunostaining.

Sections from the SNpc and STR were washed with PBS and incubated for 3 h at room temperature (RT) in PBST (PBS containing 0.1% Triton X-100) supplemented with 10% normal goat serum (NGS) for blocking and permeabilization. Sections were subsequently incubated overnight at 4°C with one of two primary-antibody combinations diluted in PBST containing 2% NGS: mouse anti-GFAP (1:250; Novus Biologicals) together with chicken anti-TH (1:300; Abcam), or mouse anti-GFAP (1:250) together with rabbit anti-pan-calcineurin A (1:250; Cell Signaling Technology).

Following three washes in PBST, sections were incubated for 1 h at RT with the appropriate fluorophore-conjugated secondary antibodies diluted in PBST containing 2% NGS. For GFAP/TH co-staining, anti-mouse Alexa Fluor 488 and anti-chicken Alexa Fluor 568 were respectively used, whereas anti-mouse Alexa Fluor 488 and anti-rabbit Alexa Fluor 647 were used for GFAP/calcineurin co-staining respectively (1:200 for each secondary antibody). Sections were then washed three additional times with PBST and mounted using ProLong Gold Antifade reagent containing DAPI (Invitrogen).

Fluorescence images were acquired using a Leica SP8 STED confocal microscope. For reconstruction of the entire SNpc region, overlapping 20× images from WT and MPTP-treated animals were acquired and digitally stitched using Adobe Photoshop v26.

### 2.3. Rat primary astrocyte culture

Astrocytes were isolated from cortical tissue following previously published methods (Garwood et al. 2011; Saha and Biswas 2015). Whole brains were harvested from 0–1-day-old Sprague-Dawley pups, the cortex was dissected and digested with trypsin for 30 min at 37°C. The digested tissue was triturated in DMEM (Gibco) containing 10% heat-inactivated FBS (Gibco), then filtered through nylon mesh to eliminate clumps. The resulting single-cell suspension was plated onto PDL-coated dishes (0.1 mg/ml working concentration) and left for 2–3 min to allow preferential adhesion of neurons. Non-adherent cells were then collected and pelleted by centrifugation (500g, 5 min), resuspended in fresh DMEM with 10% heat-inactivated FBS, and plated at either 1.2 million cells per 35 mm dish or 0.4 million cells per well of a 24-well plate. Cultures were maintained for 14 days *in vitro* (DIV), with medium replaced on alternate days.

### 2.4. Secondary cell line culture

The human astrocytoma line 1321N1 was grown in DMEM supplemented with 10% heat-inactivated FBS. The human neuroblastoma line SH-SY5Y (NCCS, Pune) was cultured in DMEM + heat-inactivated 10% FBS and differentiated over 5–7 days by adding 10 μM all-trans retinoic acid (ATRA; Sigma). Rat pheochromocytoma (PC12) cells were maintained in DMEM containing 10% heat-inactivated horse serum (HS; Gibco) and 5% heat-inactivated FBS. They were differentiated for 5–7 days in DMEM supplemented with 1% HS and 50 ng/mL nerve growth factor (NGF-β; Sigma).

### 2.5. Neurotoxin treatment

Rotenone (Sigma) was prepared in DMSO and applied to cultured cells at a final concentration of 100 nM or 200 nM. 6-Hydroxydopamine (6-OHDA; Sigma) was dissolved directly in culture medium and used at a final concentration of 100 μM. 1-Methyl-4-phenylpyridinium (MPP+) was dissolved in culture medium and added at a final concentration of 500 μM.

### 2.6. MTT cell viability assay

MTT reagent (Sigma), dissolved in culture medium, was added to cells at a final concentration of 0.5 mg/ml, followed by a 3–4 h incubation at 37°C in a humidified 5% CO2 incubator until purple formazan crystals had formed. The medium was then removed and replaced with DMSO to dissolve the crystals, with gentle agitation on an orbital shaker for 15–20 min. Absorbance was read at 570 nm.

### 2.7. Calcineurin activity assay

Cultured astrocytes were lysed on ice using a protease-inhibitor-containing lysis buffer. CN activity was measured with the Calcineurin Cellular Activity Assay Kit (Merck) per the manufacturer’s instructions. The assay relies on an RII phosphopeptide substrate specific to CN. The released free phosphate is quantified colorimetrically via the malachite green reaction at 620nm. Activity values were normalized and expressed relative to control samples (Moon et al. 2021).

### 2.8. Caspase-3 colorimetric assay

Caspase-3 enzymatic activity was measured using a commercial colorimetric assay kit (Sigma-Aldrich) according to the manufacturer’s instructions. Briefly, active caspase-3 cleavage of the specific peptide substrate releases free *p*-nitroaniline (pNA), which was quantified spectrophotometrically at 405 nm using a multi-well plate reader (Thermo Scientific). Relative fold changes in caspase-3 activity were calculated from the resulting absorbance values.

### 2.9. Intracellular Ca^2+^ imaging with Fluo-4AM

A 1 mM stock of the calcium-sensitive dye Fluo-4AM was prepared in DMSO and diluted in culture medium to a working concentration of 2 μM. Intracellular calcium levels were assessed with Fluo-4AM (Invitrogen) as per the manufacturer’s protocol: astrocytes were incubated with the dye solution for 30 min at 37°C in the dark, and fluorescence was measured at 506 nm using a multi-plate reader (Thermo Scientific).

### 2.10. Immunoblotting

Cells were lysed in RIPA buffer (Thermo Scientific) containing ProteoGuard EDTA-free protease inhibitor cocktail (Takara). Either 25 or 50 μg of protein per sample was separated by SDS-PAGE and electro-transferred onto PVDF membranes (GE Healthcare) at 100 V for 1–2 h at 4°C. Membranes were blocked for 1 h at RT in 5% BSA, then incubated overnight at 4°C with primary antibodies (see *Supplementary* Table ST3) diluted in blocking buffer. This was followed by a 1-h room-temperature incubation with appropriate HRP-conjugated secondary antibodies. Bands were visualized with Clarity Max Western ECL substrate (Bio-Rad) according to the manufacturer’s directions, and blot images were captured on an iBright imaging system (Thermo Fisher).

### 2.11. Human blood sample collection and plasma separation

Blood was drawn from Parkinson’s disease patients (n = 25) and age-matched controls (n = 20) at the Bangur Institute of Neurosciences (BIN), Kolkata, India, into EDTA vacutainers (BD Biosciences); details of the participants are provided in *Supplementary* Table ST4. Blood samples were centrifuged at 2,000 × g for 10 min at 4°C and the upper plasma fraction was collected.

### 2.12. Exosome isolation

#### 2.12.1. From astrocyte-conditioned medium (ACM)

Exosomes were purified using the miRCURY Exosome Isolation Kit for Cell/Urine/CSF (Qiagen), following the manufacturer’s protocol. Briefly, samples were first passed through a 0.2–0.22 μm syringe filter to eliminate microvesicles, and the filtrate was centrifuged at 3000×g for 10 min to remove residual cell debris. Precipitation buffer was then added and samples were left at 4°C overnight, followed by centrifugation at 3200×g for 30 min at 20°C. The resulting pellet was resuspended in resuspension buffer to yield purified exosomes for downstream analysis, or alternatively resuspended directly in the relevant cell culture medium for application to recipient cells.

#### 2.12.2. From human plasma

Plasma exosomes were isolated with the miRCURY Exosome Serum/Plasma Kit (Qiagen) according to the manufacturer’s instructions. Briefly, plasma samples were treated with thrombin for 5 min at RT and then centrifuged at 10,000×g for 5 min. Precipitation buffer was added to the resulting supernatant, which was incubated overnight at 4°C and subsequently centrifuged at 500×g for 5 min at 20°C. The pellet was resuspended in resuspension buffer to obtain purified exosomes which were subjected to downstream analysis.

### 2.13. Nanoparticle tracking analysis

Purified exosome preparations were diluted 10-fold in 1X PBS and characterized by nanoparticle tracking analysis (NTA) on a NanoSight NS300 system, following the manufacturer’s guidelines.

### 2.14. GW4869 treatment

GW4869 (Calbiochem) was dissolved in DMSO and diluted into culture medium to a final working concentration of 10 μM (Deng et al. 2022).

### 2.15. RNA extraction and quantitative real-time PCR (RT-qPCR)

Total RNA was extracted using TRIzol Reagent (Thermo Fisher) via standard phenol-chloroform extraction. For exosome-derived RNA samples, synthetic *Caenorhabditis elegans* miR-39 (cel-miR-39) was spiked into each sample prior to extraction (Bai et al. 2017) to normalize for the miRNA content analysis downstream.

miRNA cDNA synthesis was carried out using the TaqMan MicroRNA Reverse Transcription Kit (Thermo Fisher), followed by RT-qPCR with TaqMan Universal PCR Master Mix (Thermo Fisher) and the corresponding miRNA-specific TaqMan assays (Thermo Fisher; listed in *Supplementary* Table ST1). For mRNA analysis, cDNA was generated with the PrimeScript 1st Strand cDNA Synthesis Kit (Takara Bio), and RT-qPCR was performed using SYBR Green PCR Master Mix (Thermo Fisher) with the primer sequences listed in *Supplementary* Table ST2. U6 snRNA served as the endogenous control for miRNA quantification, and GAPDH mRNA served as the endogenous control for mRNA quantification; cel-miR-39-3p was used as the spike-in normalization control for exosomal samples. All reactions were run on a StepOnePlus Real-Time PCR System (Thermo Fisher), and relative expression was calculated using the comparative CT (2^-ΔΔCT) method.

### 2.16. miRNA mimic transfection

SH-SY5Y or PC12 cells were seeded and allowed to differentiate for 2 days in ATRA- or NGF-β-containing medium, respectively, prior to transfection. On the second day of differentiation, cells were transfected with either a miRNA mimic or a negative control mimic (Ambion; see *Supplementary* Table ST1) in Opti-MEM (Thermo Fisher) using Lipofectamine RNAiMAX (Invitrogen), according to the manufacturer’s protocol. Opti-MEM was replaced with the appropriate differentiation medium 6 h after transfection. Cells were cultured for a further 48 h and then exposed to the relevant neurotoxin on day 5 of differentiation.

### 2.17. Prediction of miRNA targets

Putative miRNA binding sites within the 3′UTRs of target transcripts were identified *in silico* using the target-prediction databases like TargetScan, and miRDB.

### 2.18. Cloning of the 3′UTR

To produce reporter plasmids containing 3′UTR of human NOXA, sequences were cloned into psiCHECK-2 plasmid (Promega), downstream of the Renilla luciferase coding region. The primers (Roufayel et al. 2014) used to amplify 3’UTR of human NOXA from genomic DNA are indicated in *Supplementary* Table ST2.

### 2.19. Luciferase reporter assay

SH-SY5Y cells were co-transfected with 150 ng of the psiCHECK-2 reporter construct — with or without the NOXA 3′UTR insert — together with 50 nM of either negative control mimic or miR-23a mimic, using Lipofectamine 3000 (Thermo Scientific) as per the manufacturer’s instructions. 48 h post-transfection, luciferase activity was quantified with the Dual-Glo Luciferase Assay System (Promega) on a multimode plate reader (PerkinElmer). For each construct, reporter activity was normalized as the Renilla-to-firely luciferase ratio and compared between conditions.

### 2.20. Ethics Statements

#### 2.20.1. Human Ethics

All research protocols involving human participants were reviewed and approved by the Ethical Committee of Human Subjects of CSIR-IICB. Written informed consent was obtained from all participants prior to study enrolment.

#### 2.20.2. Animal Ethics and Housing

Adult male C57BL/6 mice (8–10 weeks old, 25–30 g) were housed under controlled environmental conditions (22 ± 2 °C, 60 ± 5% relative humidity, 12-hour light/dark cycle) with ad libitum access to food and water in the animal house facility of CSIR-IICB, India. Animal protocols complied with the national guidelines set by the Committee for Control and Supervision of Experiments on Animals (CCSEA), Department of Animal Husbandry and Dairying (DAHD), Ministry of Fisheries, Animal Husbandry and Dairying, Govt. of India, and received formal approval from the Institutional Animal Ethics Committee (IAEC) of CSIR-IICB.

### 2.21. Statistical analysis

Data plots, statistical tests, and image quantification were performed using GraphPad Prism 10.0, ImageJ, and Fiji. Comparisons across more than two groups were analyzed by one-way ANOVA with Tukey’s post hoc correction, while two-group comparisons used a two-tailed, unpaired t-test with Welch’s correction. Statistical significance was set at p < 0.05 denoted *, with p < 0.01 denoted ** and p < 0.001 denoted ***. Each experiment was independently repeated at least three times unless stated otherwise, and error bars represent the mean ± SEM.

## 3. Results

### 3.1. Elevated CN expression is associated with increased DA neurodegeneration and astrogliosis in the STR of MPTP mouse model of PD

As the tissue-level expression of CN in the PD brain has not previously been reported, we sought to examine CN protein expression in the STR of the MPTP mouse model of PD. 1-Methyl-4-phenyl-1,2,3,6-tetrahydropyridine or MPTP is a neurotoxin well established to induce severe Parkinsonian pathology in humans (Langston et al. 1983; Langston 2017) as well as in rodents (Haobam et al. 2005; Muralikrishnan and Mohanakumar 1998). To generate an acute PD model, mice were administered with MPTP twice, at a gap of 16h, and sacrificed on day 8 (Mondal et al. 2023). Although the primary neuropathological hallmark of PD is degeneration of DA neurons in the SNpc, the resulting loss of nigrostriatal dopamine also adversely affects the structure and function of striatal neurons, including loss of dendritic spines (Beck et al. 2018; Stephens et al. 2005; Villalba and Smith 2010). Astrogliosis, implicated in various NDDs and neuroinflammatory conditions, is characterized by hypertrophy of astrocytic somata and processes, along with increased expression of glial fibrillary acidic protein (GFAP) (Pekny and Nilsson 2005; Sofroniew 2009; Sofroniew and Vinters 2010). We therefore performed immunohistochemistry on brain sections for tyrosine hydroxylase (TH; a DA neuron marker), GFAP (an astrocyte marker), and CN.

Substantial loss of TH positive DA neurons was observed in the SNpc of MPTP-treated mice as compared to WT (*Supplementary* Fig. S1). This loss of nigral DA neurons resulted in extensive degeneration of their striatal neurites, as evidenced by markedly reduced TH expression in the STR (Fig.1A *bottom panel*), whereas the STR of WT mice retained dense TH-positive neurites (Fig. 1A *top panel*). Notably, a corresponding increase in GFAP-expressing astroglial cells was observed in the STR of MPTP-treated mice (Fig. 1A & 1B *bottom panel*), consistent with previous reports of striatal astrogliosis in other PD models (Innamorato et al. 2010; Liu et al. 2025; Rojo et al. 2010; Viana et al. 2016). By contrast, only sparse GFAP-positive tubular astroglial structures were observed in the STR of WT mice (Fig. 1A & 1B *top panel*). Next, we sought to determine the expression of CN in the STR. An increased overall CN expression was observed in the STR of MPTP-treated mice (Fig. 1B *bottom panel*) as compared to that of WT (Fig. 1B *top panel*). Quantification of fluorescence intensities confirmed a significant decrease in TH expression alongside significant increases in GFAP and CN expression in the STR of MPTP-treated mice relative to WT, as shown in Fig. 1C–1E, respectively.

**Figure 1:**
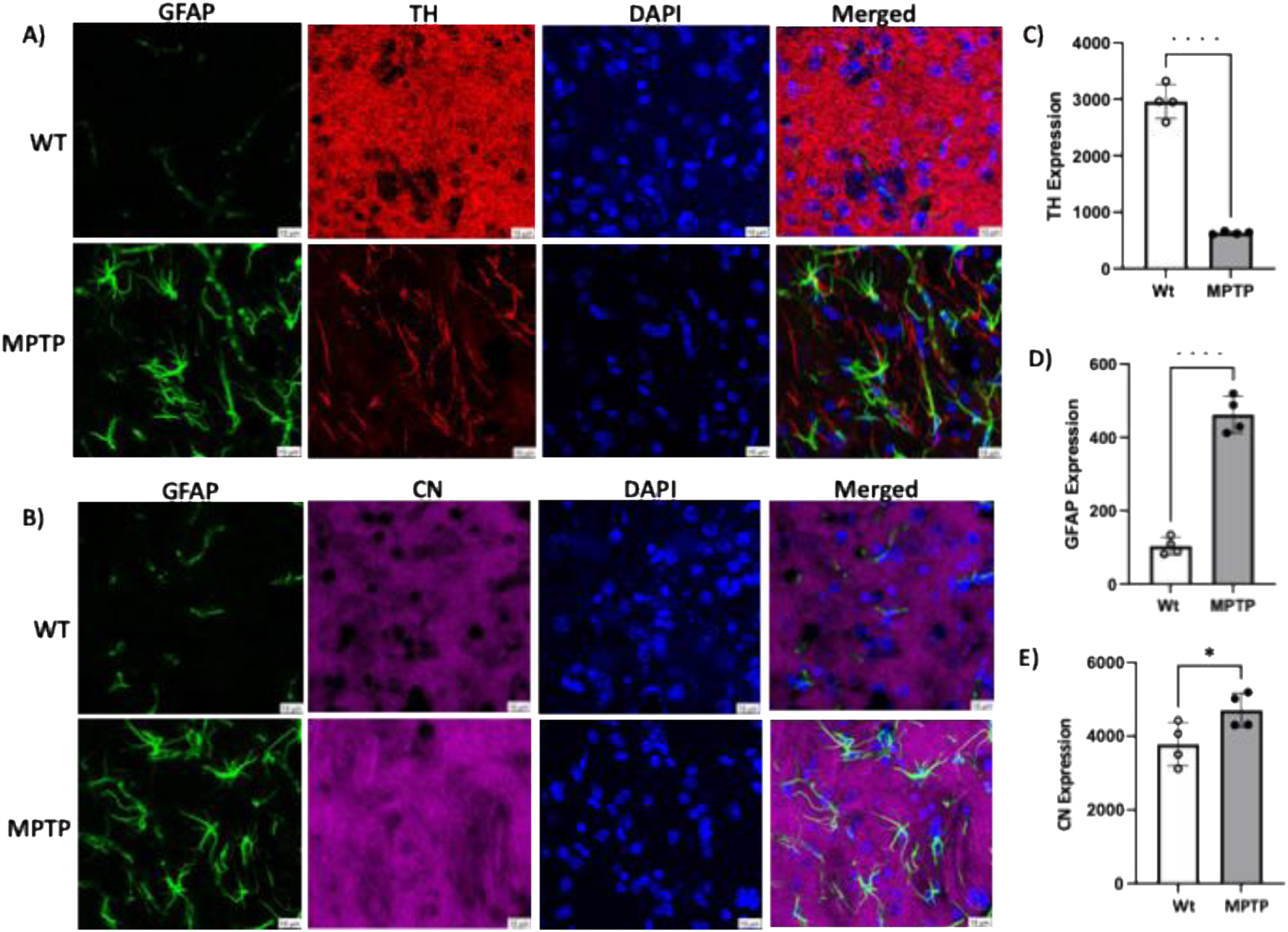
Increased CN expression and astrogliosis in the STR of MPTP mice: A & B) Immunohistochemistry of STR sections from WT and MPTP-treated mice, imaged under confocal microscope (Leica Sp8 STED microscope). GFAP (green), TH (red), CN expression (magenta), DAPI (blue) and merged images. C) D) and E) Fluorescence intensities were measured by using ImageJ software. Bar diagrams represent corresponding pixel densities for the TH, GFAP and CN expression levels. Results are plotted as Mean ± S.E.M where *p < 0.05 and ****p < 0.0001 (3 images per animal and n = 4 per group).

Therefore, our results suggest that increased CN expression accompanies DA denervation and astrogliosis in the STR of MPTP-treated mice.

### 3.2. Rotenone induces Ca²⁺ elevation and CN activation in astrocytes

Having observed increased CN protein expression with accompanying astrogliosis in the STR of the MPTP mouse model of PD, we sought to investigate the consequences of CN expression at the intracellular level, particularly in astrocytes, since it remains unexplored till date. To establish a cellular model of PD, cultured astrocytes were treated with Rotenone, a mitochondrial complex I inhibitor widely used to model PD-associated neurotoxic stress (Innos and Hickey 2021). Rotenone was selected over MPP+ (the active metabolite of MPTP) because it readily crosses the plasma membrane of both astrocytes and neurons, whereas MPP+ uptake depends on the dopamine transporter (DAT), which is present in neurons (Javitch et al. 1985; Richardson et al. 2005).

Human astrocytoma cells (1321N1) (Banerjee et al. 2013) were treated with varying doses of Rotenone for 24 h followed by MTT assay which showed approximately 55-65% survival at 200nM dose (Fig. 2A). Subsequently, cells were treated with 200 nM Rotenone for varying durations, and CN-A expression was assessed by immunoblotting. We used an antibody directed against the N-terminus (Novus Biologicals), which can detect both the full-length (FL, 59-61 kDa) and truncated (45-48 kDa) forms of CN-A (Fig. 2B). Relative to FL CN-A, truncated CN-A levels increased by approximately 2-fold and 3-fold at 4 h and 8 h of Rotenone treatment respectively, as compared with untreated controls (Fig. 2C).

**Figure 2:**
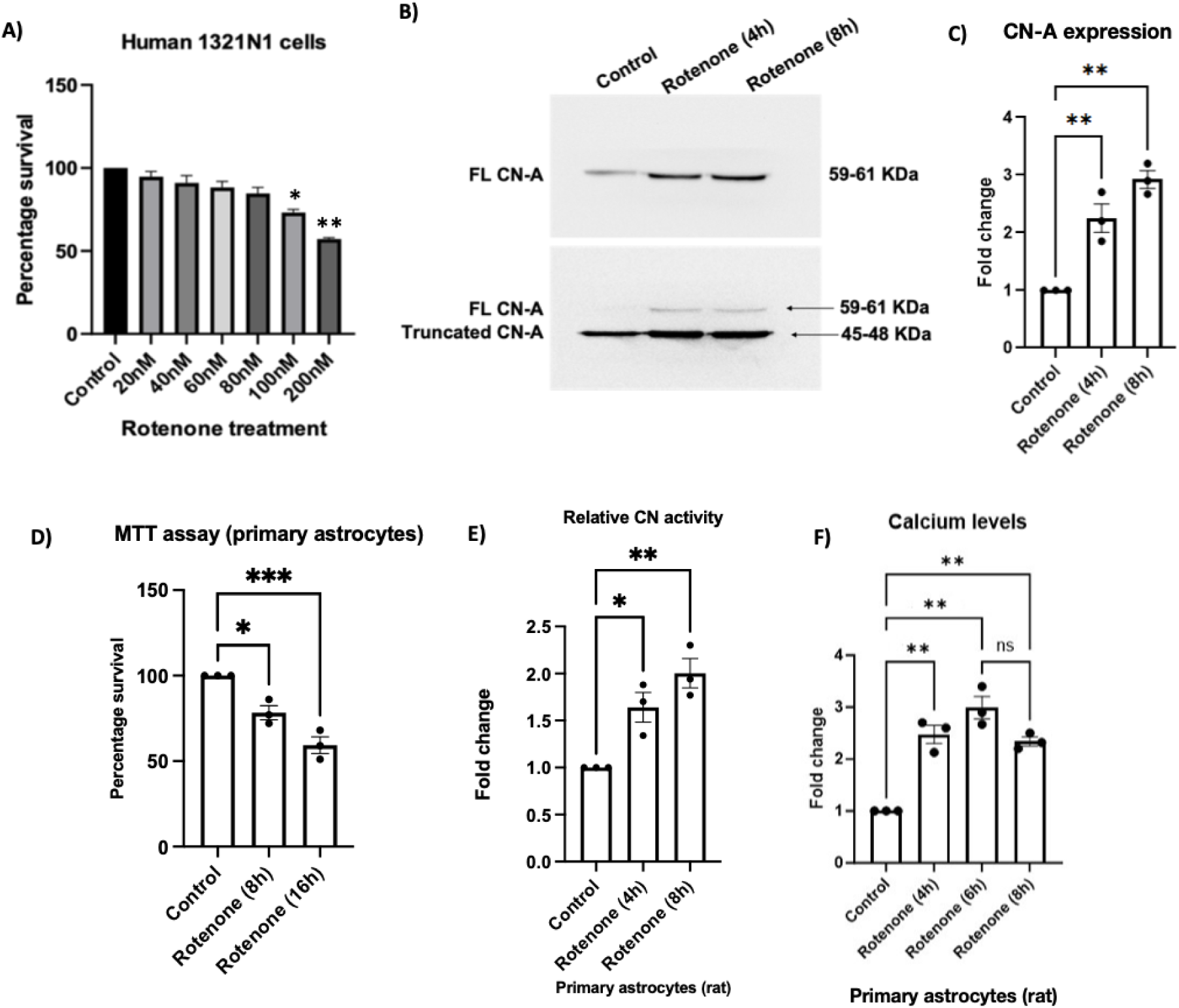
Rotenone-induced activation of Ca²⁺/CN in astrocytes. A) Human astrocytoma (1321N1) cells were treated with Rotenone (20, 40, 60, 80, 100, and 200nM) for 24 h, and cell viability was assessed by MTT assay and expressed as a percentage relative to untreated control cells taken as 100% (n = 4). B) 1321N1 cells were left untreated (control) or treated with 200nM Rotenone for the indicated time-points, and total protein lysates were subjected to immunoblotting for CN-A using an antibody against its N-terminus. C) Densitometric quantification of truncated CN-A levels, normalized to FL CN-A (n = 3) as depicted in (B). D) Primary rat astrocytes (14 DIV) were treated with 100nM Rotenone for 0h (control), 8h, or 16h and cell viability was assessed by MTT assay and expressed as a percentage relative to untreated control taken as 100% (n = 3). E) Control and Rotenone-treated (100 nM; 4h, 8h) primary rat astrocyte lysates were incubated with an RII phosphopeptide substrate, and CN enzymatic activity was assessed by colorimetric detection of free phosphate release at 620 nm. Results are expressed as relative fold change with respect to control (n = 3). F) Control and Rotenone-treated (100 nM; 4h, 6h, 8h) primary rat astrocytes were loaded with Fluo-4AM and fluorometric readings were obtained at 506 nm (n = 3). Asterisks denote statistically significant differences and “ns” denotes non-significant differences relative to control, unless otherwise indicated. Data are presented as mean ± SEM; *p < 0.05, **p < 0.01, ***p < 0.001.

To extend these findings to a physiologically relevant model, primary astrocytes (14 DIV) were cultured from neonatal rat brains and treated with 100 nM Rotenone (Rathinam et al. 2012) over a time course of 0– 16 h. Cell viability, assessed by MTT assay, showed approximately 75% and 60% cell survivability at respectively 8 h and 16 h of treatment relative to untreated cells (Fig. 2D). To check CN activity at the enzymatic level, a CN enzyme assay was performed in rat primary astrocytes using a commercial kit (Merck) which revealed increased CN activity at 4 h and 8 h of Rotenone treatment as compared to untreated (control) cells (Fig. 2E). As CN is a Ca²⁺-dependent protein phosphatase requiring Ca²⁺ binding to its catalytic subunit for maximal activity, intracellular Ca²⁺ levels were also examined. Rat primary astrocytes were loaded with the cell-permeable calcium indicator Fluo-4AM, and fluorescence was measured at Ex/Em 494/506 nm. As compared to control (untreated) cells, a significant increase in fluorescence intensity, indicating Ca²⁺ surge, was observed in primary astrocytes at 4 h and 6 h of Rotenone treatment, which was sustained through 8 h (Fig. 2F).

Collectively, these results demonstrate a concurrent increase in intracellular CN expression and its activity in astrocytes, during the early stages of Rotenone treatment.

### 3.3. Rotenone-induced decrease in miR-23a expression within astrocytes

Having established that Rotenone induces Ca²⁺ elevation and CN activation in astrocytes, we next examined whether miR-23a, a reported downstream target of CN signalling, is altered under these conditions. As mentioned earlier, (Lin et al. 2009) has reported that CN pathway activation can directly upregulate miR-23a expression in cardiomyocytes. Notably, within brain tissue, miR-23a is an astrocyte-enriched miRNA (Smirnova et al. 2005) that has also been reported to be upregulated in the serum of patients with PD and other neurodegenerative conditions (Barbagallo et al. 2020). However, the mechanistic role of miR-23a in astrocytes remains unreported to date. Given this, astrocyte-enriched miR-23a emerged as a compelling candidate for investigation downstream of CN activation, particularly in the context of PD pathogenesis.

To this end, 1321N1 cells were treated with Rotenone at the previously determined dose of 200 nM for varying durations, and intracellular miR-23a levels were quantified by RT-qPCR. A significant decrease in miR-23a expression was observed at 4h and 8h of treatment (Fig. 3A), coinciding with the same time-window in which CN activation was observed earlier. Interestingly, this finding is in contrast to that reported by (Lin et al. 2009) in cardiomyocytes, where expression of a constitutively active form of CN resulted in marked upregulation of miR-23a. To further validate our observation, primary rat astrocytes were treated with Rotenone (100 nM) for 4h and 8h, and a similar time-dependent decrease in miR-23a expression was observed (Fig. 3B). Collectively, these findings demonstrate that Rotenone treatment is associated with decreased intracellular miR-23a levels in astrocytes, despite the concurrent activation of CN, in contrast to the positive regulatory relationship previously reported in cardiomyocytes. The mechanistic basis for this cell-type-specific discrepancy is explored in subsequent sections.

**Figure 3.**
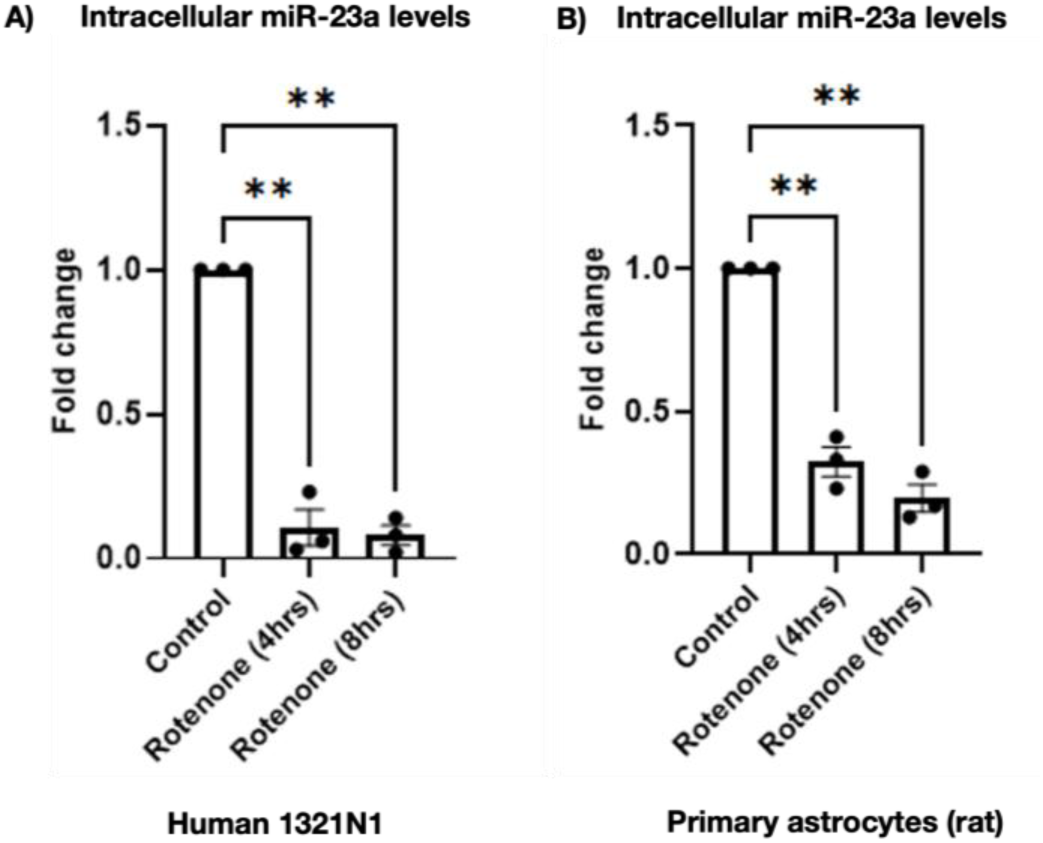
Rotenone decreases intracellular miR-23a levels in astrocytes. Human 1321N1 cells (A) and primary rat astrocytes (B) were treated with Rotenone (200 nM and 100 nM, respectively) for the indicated time points or left untreated (control). Intracellular miR-23a levels were assessed by RT-qPCR (n = 3 for each condition). Asterisks indicate statistically significant differences relative to control. Data are presented as mean ± SEM; **p < 0.01.

### 3.4. Rotenone treatment induces exosome-mediated release of miR-23a from astrocytes

Although miR-23a has been reported as a downstream target of the CN pathway, our results demonstrated that CN activation did not, in fact, increase miR-23a levels in astrocytes. To address this discrepancy, we hypothesized that astrocytes might release miR-23a into the extracellular milieu, as astrocytes can release intracellular cargo via exosomes and thereby mediate communication with neighbouring neurons (Venturini et al. 2019). We therefore investigated whether miR-23a is released from astrocytes via exosomes under our experimental conditions.

To this end, 1321N1 cells were treated with Rotenone (200nM), and astrocyte-conditioned medium (ACM) was collected at 0h (control), 4h, and 8h of treatment. Exosomes were isolated from the ACM using the miRCURY Exosome Isolation Kit (Qiagen), total exosomal RNA was isolated and miR-23a level was quantified by RT-qPCR. A significant increase in exosomal miR-23a levels were observed at 4h and 8h of Rotenone treatment relative to control (Fig. 4A), coinciding with the time-window of decreased intra-astrocytic miR-23a levels previously observed (Fig. 3A & 3B). These findings indicate that the Rotenone-induced reduction in intracellular miR-23a is accompanied by increased exosomal miR-23a levels, suggesting enhanced extracellular release of miR-23a from astrocytes.

**Figure 4:**
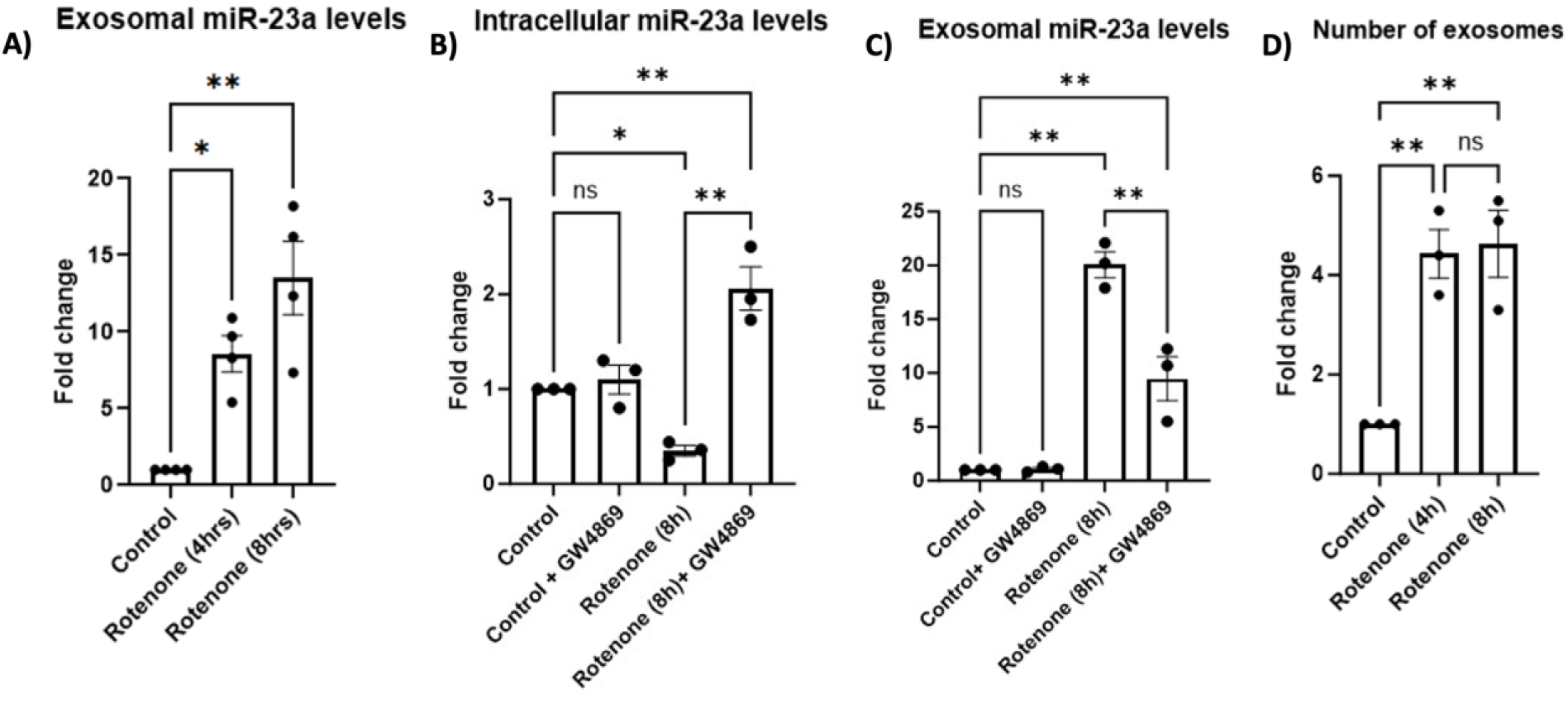
Rotenone promotes miR-23a release from astrocytes via exosomes. A) 1321N1 cells were either untreated (control) or treated with Rotenone for 4h or 8h, and ACM was collected. Exosomes were isolated from the ACM, total exosomal RNA was extracted and miR-23a levels were assessed by RT-qPCR (n = 4). B & C) 1321N1 cells were divided into four groups: (i) untreated control; (ii) GW4869 (10 μM, 24 h) alone; (iii) Rotenone alone (8 h); or (iv) GW4869 pre-treatment (10 μM, 24 h) followed by Rotenone (8 h). Expression of miR-23a in exosomes isolated from ACM (C) and intracellular miR-23a levels (B) were assessed by RT-qPCR (n = 3 for each). D) Exosomes were isolated from ACM collected under untreated (0h) and Rotenone-treated (200 nM; 4h and 8h) conditions, and total exosome number was determined by nanoparticle tracking analysis (NTA) (n = 3). Asterisks indicate statistically significant differences and “ns” indicates non-significant differences relative to control, unless otherwise indicated. Data are presented as mean ± SEM; *p < 0.05, **p < 0.01.

It is reported that exosome release is dependent on ceramide generation by neutral sphingomyelinase 2 (nSMase2), and treatment with nSMase2 inhibitors such as GW4869 can reduce exosome secretion (Kosaka et al. 2010; Mukherjee et al. 2016; Trajkovic et al. 2008). To further validate exosomal release of miR-23a by astrocytes, 1321N1 cells were pre-treated with GW4869 (10 μM) for 24h, followed by Rotenone treatment for 8h, and miR-23a levels were assessed both intracellularly and in exosomes isolated from the ACM by RT-qPCR. In the ACM, GW4869 significantly reduced the Rotenone-induced increase in exosome-derived miR-23a, supporting an exosome-dependent mechanism of miR-23a release (Fig. 4C). Conversely, inhibition of exosome release with GW4869 significantly increased intracellular miR-23a levels following Rotenone treatment (Fig. 4B), consistent with reduced extracellular release of miR-23a observed in Fig. 4C. We next sought to determine if there is any alteration in the number of exosomes released upon Rotenone treatment. Exosomes were isolated from the ACM under untreated (0h) and Rotenone-treated (200 nM; 4h and 8h) conditions, and total exosome count was determined by nanoparticle tracking analysis (NTA), following the methodology previously described in (Bhattacharyya et al. 2023). A significant increase in exosome number was observed at both 4h and 8h of Rotenone treatment, relative to untreated conditions (Fig. 4D).

Together, these findings suggest that Rotenone exposure promotes exosome-associated release of miR-23a from astrocytes, accompanied by a reduction in intracellular miR-23a levels and an overall increase in exosome release.

### 3.5. Neuroprotective function of astrocyte-derived exosomes and miR-23a

We next investigated the functional consequence of astrocyte-derived exosomes released during neurotoxic stress and examined whether these exosomes exert a protective or detrimental effect on neuronal cells. To this end, 1321N1 cells were treated with Rotenone for 8h, after which ACM was collected. Exosomes were isolated from the ACM as described above, *indicated as Exosomes (8h)*, resuspended in culture medium (Ghoshal et al. 2021), and added overnight to neuronally differentiated SH-SY5Y cells. The cells were subsequently exposed to Rotenone at 200 nM concentration for 24 h, previously reported to induce apoptosis in serum-containing SH-SY5Y cultures (Newhouse 2004), and cell viability was assessed by MTT assay. Notably, Rotenone-treated SH-SY5Y cells pre-incubated with ACM-derived exosomes exhibited significantly higher viability than cells treated with Rotenone alone (Fig. 5A). Exosome treatment alone did not significantly alter baseline cell viability in untreated cells (Control vs. Control + Exosome, ns), indicating that exosomes did not exert detectable cytotoxic effects under basal conditions. These findings indicate that exosomes released from Rotenone-treated astrocytes can confer protection against Rotenone-induced neuronal toxicity.

**Figure 5.**
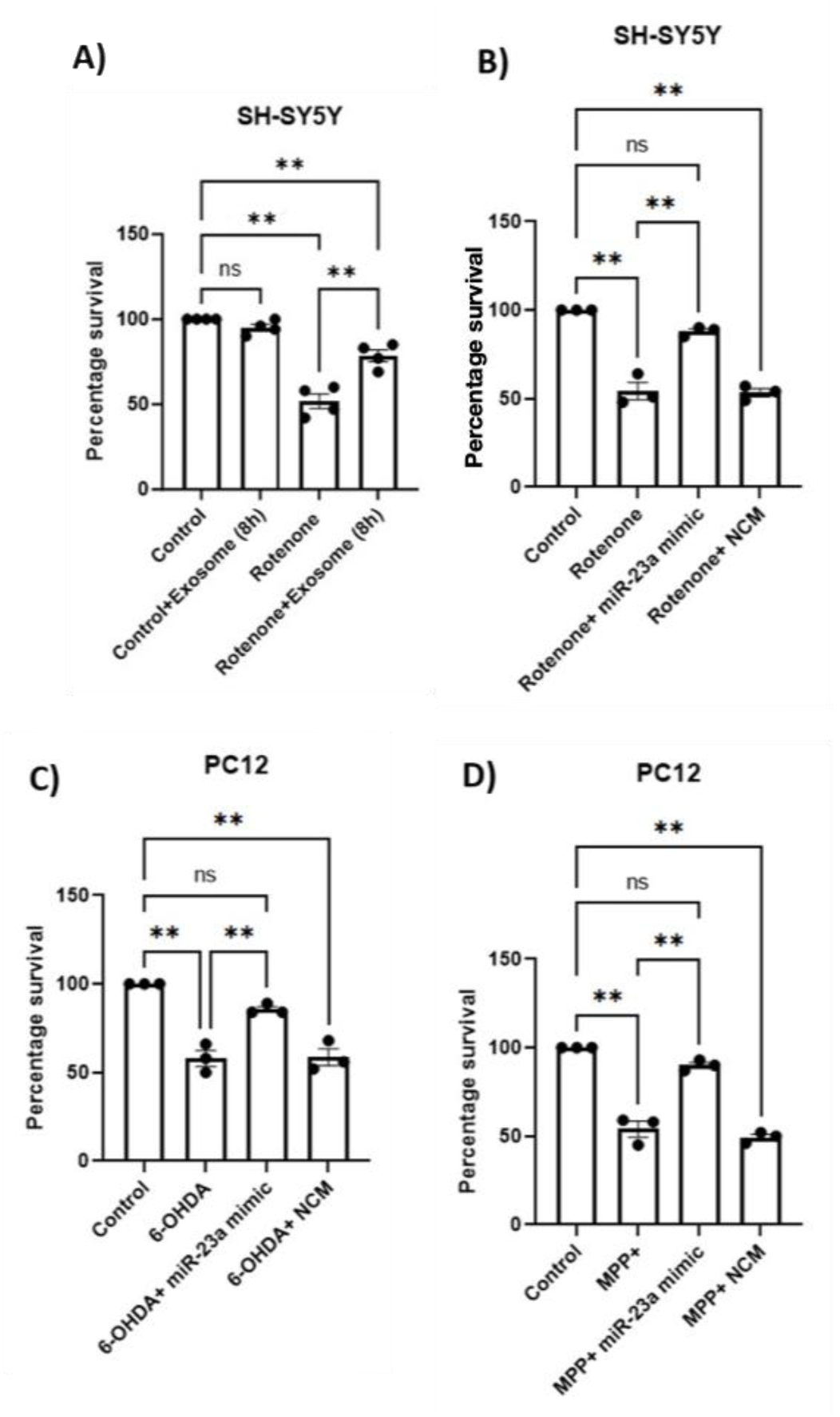
Astrocyte-derived exosomes and miR-23a protect neurons from neurotoxic stress. A) SH-SY5Y cells were divided into four groups: (i) untreated control; (ii) Rotenone alone (200 nM, 24 h); (iii) exosomes isolated from ACM (collected 8h post-Rotenone treatment), added overnight, without subsequent Rotenone treatment; or (iv) the same exosome pre-treatment followed by Rotenone treatment (200 nM, 24 h). Cell viability was assessed by MTT assay, and percentage cell viability was calculated relative to untreated control (n = 4). B) SH-SY5Y cells were either untreated (control), treated with Rotenone (200 nM, 16 h), or pre-transfected with miR-23a mimic or negative control mimic (NCM) for 48 h prior to Rotenone treatment (16 h). C) NGF-primed PC12 cells were either untreated (control), treated with 6-OHDA (100 μM, 16 h), or pre-transfected with miR-23a mimic or NCM for 48 h prior to 6-OHDA treatment (16 h). D) NGF-primed PC12 cells were either untreated (control), treated with MPP+ (500 μM, 16 h), or pre-transfected with miR-23a mimic or NCM for 48 h prior to MPP+ treatment (16 h). In (B–D), cell viability was assessed by MTT assay, and percentage cell viability was calculated relative to untreated control (n = 3 for each). Data are presented as mean ± SEM. Asterisks indicate statistically significant differences, and “ns” indicates no significant difference. **p < 0.01.

However, since exosome contains diverse molecular cargo (Lee et al. 2024), these findings alone do not establish whether miR-23a specifically contributes to the observed neuroprotective effect. We therefore directly examined the effect of miR-23a on neuronal survival. Neuronally differentiated SH-SY5Y cells were pre-transfected with a miR-23a mimic (for transient overexpression) or a negative control mimic (NCM) for 48h prior to Rotenone treatment for 16 h. MTT assay revealed that miR-23a mimic expression significantly attenuated Rotenone-induced loss of cell viability, restoring viability to a level not significantly different from untreated control (Fig. 5B).

To confirm that this neuroprotective effect was not cell line or neurotoxin-specific, the experiment was repeated in NGF-primed PC12 cells treated with 6-OHDA (100 μM) (Sanphui et al. 2020) or MPP+ (500 μM) (Wang et al. 2011). MTT assays performed 16 h after either treatment yielded similar neuroprotective results, with miR-23a mimic protecting primed PC12 cells against both 6-OHDA (Fig. 5C) and MPP+-induced (Fig. 5D) loss of cell viability. Collectively, these findings suggest that miR-23a may contribute, at least in part, to the neuroprotective effects of astrocyte-derived exosomes against neurotoxin-induced cellular stress.

### 3.6. miR-23a regulates the pro-apoptotic BH3-only protein NOXA through its 3′UTR in neurons

We next investigated the probable molecular mechanism underlying the neuroprotective effect of miR-23a. miR-23a has been shown to exert an anti-apoptotic role in several pathological contexts, including traumatic brain injury (Sabirzhanov et al. 2014) and radiation-induced CNS toxicity associated with cancer radiotherapy (Sabirzhanov et al. 2020). Moreover, we have previously reported that in 6-OHDA mediated cellular model of PD, apoptosis occurs in primed SH-SY5Y cells via induction of the pro-apoptotic molecule PUMA at as early as 4h of treatment (Bhattacharyya et al. 2023). Therefore, we sought to examine whether miR-23a overexpression affects PUMA expression levels in our cellular models of PD. The following experimental conditions were compared in primed SH-SY5Y cells: (i) untreated control, (ii) cells treated with 6-OHDA (100 μM) for 4h, (iii) cells pre-transfected with either miR-23a mimic (overexpression construct) or (iv) negative control mimic (NCM) for 48h prior to 6-OHDA treatment (4h). Transfection with the miR-23a mimic significantly reduced PUMA transcript levels (Fig. 6A), which were elevated by over 2-fold within 4h of 6-OHDA treatment. Previously, (Sabirzhanov et al. 2014) has demonstrated that miR-23a can functionally target the 3′UTRs of mouse BH3-only proteins - PUMA and NOXA. However, *in silico* analysis using miRNA target prediction tools and databases, namely TargetScan and miRDB, did not predict human PUMA as a direct target of miR-23a, rather miR-23a was predicted to bind to the 3’UTR of human NOXA. This observation was consistent with findings by (Roufayel et al. 2014), who reported direct binding of miR-23a on the 3′UTR of human NOXA.

**Figure 6:**
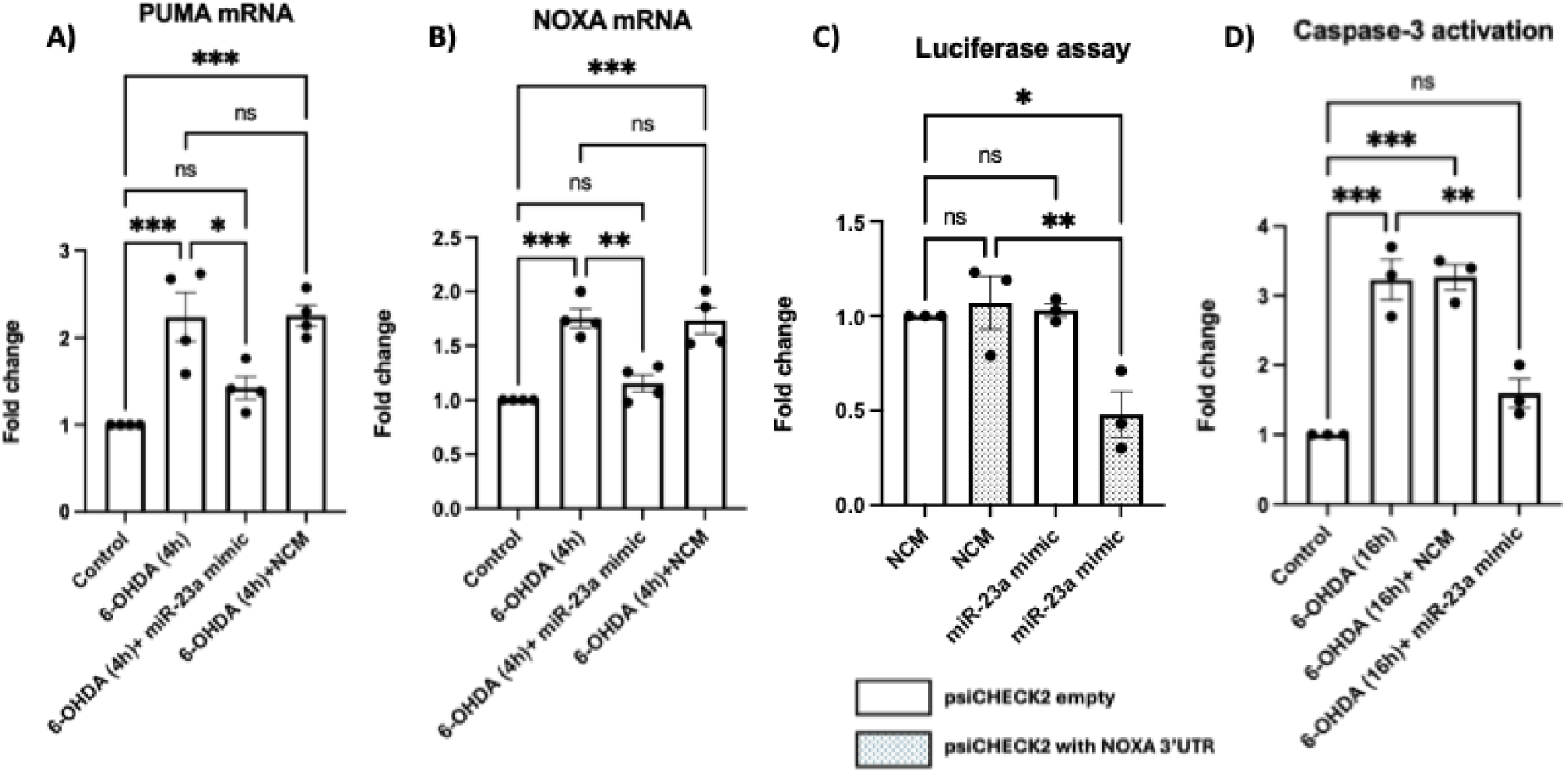
miR-23a directly regulates NOXA and suppresses pro-apoptotic signalling in neurons. A & B) Primed SH-SY5Y cells were either untreated (control), treated with 6-OHDA (100μM) for 4 h, or pre-transfected with miR-23a mimic or NCM for 48h prior to 6-OHDA treatment. Total RNA was isolated, and mRNA levels of PUMA (A) and NOXA (B) were measured by RT-qPCR, with GAPDH used as the endogenous control for normalization (n = 4). C) Primed SH-SY5Y cells were co-transfected with one of the following combinations: NCM and empty psiCHECK-2 vector; or NCM and psiCHECK-2 reporter plasmid containing the NOXA 3′UTR; or miR-23a mimic and empty psiCHECK-2 vector; or miR-23a mimic and psiCHECK-2 reporter plasmid containing the NOXA 3′UTR. Luciferase assay was performed 48 h post-transfection, with RLuc signal normalized to FLuc signal. Fold change for each condition was calculated relative to control (empty psiCHECK-2 vector + NCM), set as 1 (n = 3). D) Primed SH-SY5Y cells were either untreated (control), treated with 6-OHDA (100μM) for 16 h, or pre-transfected with miR-23a mimic or NCM for 48h prior to 6-OHDA treatment. Cell lysates were incubated with the chromophore-labeled substrate DEVD-pNA, and caspase-3 activity was measured by absorbance at 405 nm (n = 3). Asterisks indicate statistically significant differences and “ns” indicates non-significant differences relative to control, unless otherwise indicated. Data are presented as mean ± SEM; *p < 0.05, **p < 0.01, ***p < 0.001.

To test whether NOXA is implicated in our cellular model of PD, NOXA mRNA levels were assessed under the same experimental conditions described above. While 100 μM 6-OHDA rapidly increased NOXA mRNA levels within 4 hours, overexpression of miR-23a successfully mitigated this response (Fig. 6B). We next sought to confirm direct binding of miR-23a to the NOXA 3′UTR in SH-SY5Y, adapting the approach of (Roufayel et al. 2014). The NOXA 3′UTR was cloned downstream of the *Renilla* luciferase gene in the psiCHECK-2 plasmid (Promega), and a luciferase reporter assay was performed. SH-SY5Y cells were co-transfected under one of four conditions: (i) NCM with empty psiCHECK-2, (ii) NCM with psiCHECK-2 containing the NOXA 3′UTR, (iii) miR-23a mimic with empty psiCHECK-2, or (iv) miR-23a mimic with psiCHECK-2 containing the NOXA 3′UTR. At 48 h post-transfection, *Renilla* (RLuc) and firefly (FLuc) luciferase signals were measured using the Dual-Glo Luciferase Assay System (Promega), and relative luciferase activity was calculated by normalizing RLuc to FLuc. Luciferase activity was significantly reduced in cells co-transfected with the NOXA 3′UTR-containing plasmid and miR-23a mimic as compared to the cells co-transfected with the NOXA 3′UTR-containing plasmid and NCM (Fig. 6C), consistent with direct post-transcriptional regulation of NOXA through its 3′UTR by miR-23a.

Finally, as both PUMA and NOXA belong to the pro-apoptotic BH3-only protein family, we sought to determine whether miR-23a overexpression suppresses downstream apoptotic signalling. We have previously reported (Bhattacharyya et al. 2023) the activation of caspase-3 at 16h of 6-OHDA (100 μM) treatment in primed SH-SY5Y cells. Here, miR-23a overexpression was found to significantly attenuate 6-OHDA-induced caspase-3 activity in primed SH-SY5Y cells (Fig. 6D), as assessed by colorimetric assay following the method of (Bhattacharyya et al. 2023). Collectively, these results suggested that miR-23a exerts its neuroprotective effect by suppressing a pro-apoptotic BH3-only protein network, at least in part, through direct repression of NOXA and a concomitant reduction of PUMA expression, thereby limiting apoptotic signalling.

### 3.7. miR-23a expression is elevated in plasma-derived exosomes from PD patients

Since our cell culture-based results strongly indicated that miR-23a released within astrocyte-derived exosomes plays an important role in PD pathogenesis, we sought to determine whether miR-23a could be detected in exosomes isolated from clinical PD samples. We have previously reported the presence of exosomes in PD patient plasma and determined their size and number by nanoparticle tracking analysis (NTA) (Bhattacharyya et al. 2023). Using the same pool of PD patient samples and age-matched controls, we now determined the expression of miR-23a in plasma-derived exosomes, following the methodology described in (Bhattacharyya et al. 2023). Briefly, blood samples were collected from PD patients (> 40 years) along with age-matched controls without any history of neurological disorder. Plasma was separated by centrifugation, and intact exosomes were isolated using the miRCURY Exosome Isolation Kit according to the manufacturer’s protocol (Helwa et al. 2017; López-Pérez et al. 2021). Total RNA was subsequently isolated from the exosomes, and RT-qPCR was performed to assess miR-23a expression in both PD patient and non-patient (control) samples. Our results showed that miR-23a levels were significantly higher in plasma-derived exosomes from patients with PD than in age-matched controls (Fig. 7). Thus, these findings support the clinical relevance of our observations in cellular models of PD-associated neurotoxic stress.

**Figure 7:**
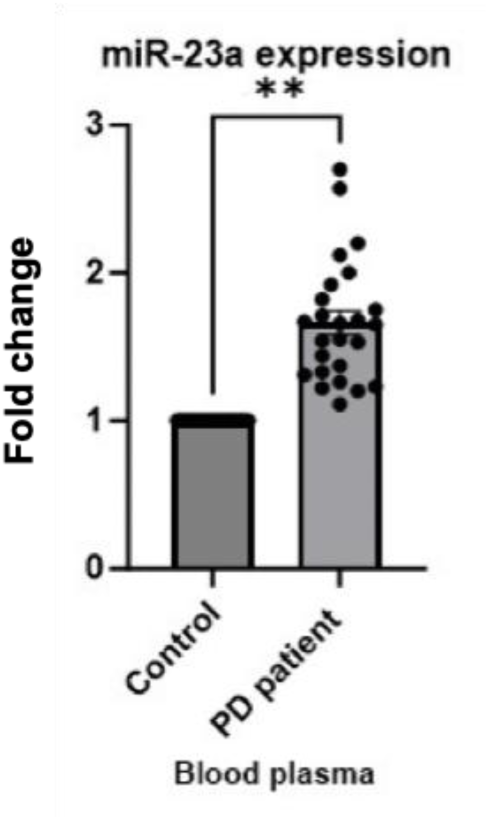
miR-23a expression is increased in exosomes derived from human PD patient plasma. Exosomal miR-23a expression was measured in plasma samples from PD patients (age > 40 years; n = 25) compared with age-matched controls (n = 20). Asterisks indicate statistically significant differences relative to control. Data are presented as mean ± SEM; **p < 0.01.

## 4. Discussion

Our study identifies astrocytic Ca²⁺/CN upregulation as a previously unrecognized component of the cellular response to PD-associated neurotoxic stress. We first observed increased CN protein expression in the STR of MPTP-treated mice, concomitant with loss of striatal TH-positive innervation and increased GFAP-positive astroglial reactivity. Our findings complement previous evidence of enhanced CN enzyme activity in the STR of MPTP-treated mice and its contribution to PD-associated behavioural and structural abnormalities in neurons (*Supplementary data*, Mondal et al. 2023). On the other hand, (Tamburrino et al. 2015) showed (by immunoblotting) that in MPTP mice, NFATc3, a downstream target of CN signalling, and astrocyte marker GFAP are increased in the STR which was attenuated by pharmacological CN inhibition. These findings strengthen the importance of activated CN in MPTP mouse models of PD. Previously, (Zhuang et al. 2020) had reported that a substantial increase in intracellular Ca²⁺/CN signalling occurs in neuronal SH-SY5Y cells following 6-OHDA/ascorbate treatment. On the other hand, (Luo et al. 2014) showed that in transgenic mice, expression of the PD-related human α-synuclein A53T mutation promoted CN activity in midbrain DA neurons while pharmacological inhibition of CN reduced the α-synuclein-associated DA neuron loss. Together with our observations, these studies indicate that activated CN signalling represents a recurrent feature of experimental PD models, independent of the type of neurotoxin or transgene expression used. However, most previous investigations have focused on CN activity in neurons. Our study overcomes this lacuna by elucidating for the first time at cellular levels, the molecular significance of CN activation in non-neuronal cells (astrocytes) during PD progression.

We used human secondary astrocytoma cell line 1321N1 and primary astrocyte culture from neonatal rat brain, to validate the intracellular activation of Ca²^+^/CN in astrocytes, under the influence of Rotenone. Rotenone induced a strong increase in intracellular Ca²⁺ accompanied by increased CN-A expression and enzymatic activity within the astrocytes. These changes occurred at early time points (4h-8h), preceding substantial loss of astrocyte viability, suggesting that Ca²⁺/CN activation represents an early cellular response to Rotenone rather than simply a consequence of extensive cellular injury. Interestingly, under physiological conditions, CN is predominantly expressed in neurons, while its expression in astrocytes is reported to be low or undetectable (Dawson et al. 1994; Goto et al. 1986; Norris et al. 2005). In pathological context, most studies on Ca²⁺/CN signalling in astrocytes, have been performed in another major NDD like Alzheimer’s disease (AD), traumatic brain injury or inflammatory models, rather than PD (Sompol and Norris 2018; Lim et al. 2023). For instance, a seminal paper by (Norris et al. 2005) demonstrated that CN activation can drive reactive/inflammatory astrocyte phenotypes and that CN signalling is increased in aging and AD models. Similarly, (Pyrzynska et al. 2001) reported expression of the CN-A subunit in reactive astrocytes and that inhibition of CN can affect the survival/proliferation of these astrocytes. More recently, (Sompol et al. 2017) reported that a highly active CN phosphatase fragment and NFAT4 appear in astrocytes in direct proportion to astrocyte activation, driving glutamate dysregulation and neuronal hyperexcitability in mouse model of AD. Thus, although Ca²⁺/CN signalling has been extensively characterized in reactive astrocytes in AD, its activation and downstream consequences in astrocytes under PD-associated neurotoxic stress remain poorly characterized.

In order to assess the significance of Ca²⁺/CN activation in astrocytes, we decided to explore the expression level of an astrocyte-enriched miRNA, miR-23a (Smirnova et al. 2005), since (Barbagallo et al. 2020) had shown that miR-23a was among the six miRNAs significantly increased in PD patient serum. Moreover, (Lin et al. 2009) had shown that constitutively active CN could increase miR-23a expression, whereas CN inhibition suppressed this increase of miR-23a in cardiomyocytes. In our astrocyte cultures, Rotenone treatment resulted in a significant decrease in miR-23a levels within 4h-8h. This result seemed inconsistent with that reported by (Lin et al. 2009), although the divergence may reflect differences in cell type, cellular context or the duration and nature of the stimulus. Moreover, (Lin et al. 2009) measured steady-state intracellular miR-23a levels following sustained CN activation, without assessing extracellular/exosomal export. Interestingly, (Hudson et al. 2014) had reported that while miR-23a expression is upregulated downstream of Ca²⁺/CN activation, these miR-23a are increasingly packaged out of skeletal muscle cells via exosomes. Their findings prompted us to probe if miR-23a is exported via exosomes in our model. We observed an increase in miR-23a expression in the ACM-derived exosomes, post-Rotenone treatment.

Conversely, inhibition of exosome biogenesis with GW4869 resulted in an increase in intracellular miR-23a levels. Thus, our data do not exclude the possibility that CN activation promotes miR-23a transcription in astrocytes; rather, they suggest that any such increase in production is outpaced by a concurrent, cellular stress-mediated enhancement of exosomal export, such that the net intracellular pool of miR-23a declines even as total cellular miR-23a output may in fact be elevated. We also observed an overall increase in exosome secretion upon Rotenone insult, when compared to untreated astrocytes. These findings are consistent with previous reports that suggest that Rotenone induces exosome release in cancer stem cells (Kumar et al. 2015) while (Pan-Montojo et al. 2012) demonstrated that Rotenone increased the number of exosomes and promoted exocytosis of α-synuclein in enteric neurons from mice. Although direct involvement of CN signalling for exosome biogenesis is not yet reported, evidences of calcium influx to activate intracellular signalling pathways for enhanced exosome secretion has been shown (Savina et al. 2003). However, increased exosome secretion or production is not Rotenone-specific and other PD models have also recapitulated increased exosome release from astrocytes. For instance, (Wang et al. 2023) found that the number of astrocyte-derived EVs significantly increased following A53T α-synuclein expression or exposure to α-syn aggregates on primary mouse astrocytes. This was accompanied by increased Alix and multivesicular bodies which are involved in exosome biogenesis. Another important report by (Leggio et al. 2022) showed increased release of small EVs from ventral midbrain and striatal astrocytes-regions most affected during PD. So far, our findings indicate that neurotoxin-associated Ca²⁺/CN activation coincides with increased release of miR-23a in exosomes derived from astrocytes.

Next, we sought to investigate whether the altered release of astrocytic exosome-associated miR-23a has any functional consequences for neurons. Exosomes are well established as important mediators of astrocyte-neuron crosstalk under both physiological (Men et al. 2019) and stress conditions (Venturini et al. 2019). In the CNS, exosomes released by astrocytes and microglia predominantly support homeostatic functions and play important roles in regulating neuroinflammation (Delpech et al. 2019; Dickens et al. 2017). In our cellular models of PD, we found that astrocyte-derived exosomes were neuroprotective. This result was in concordance with that of (Leggio et al. 2022) which showed that small EVs from ventral midbrain and nigrostriatal astrocytes can counteract H2O2-induced caspase-3 activation and MPP+-induced mitochondrial dysfunction in differentiated SH-SY5Y cells. Another report by (De Rus Jacquet et al. 2021) showed altered EV biogenesis in astrocytes generated from induced pluripotent stem cells from PD patients with LRRK2 mutation; the EVs were internalized by DA neurons, leading to dysfunctional astrocyte-neuron cross-talk. Now, exosomal cargo can be diverse, so we decided to pinpoint whether miR-23a could account, at least in part, for this neuroprotective activity of exosomes. Although context dependent, miR-23a has been implicated in regulating different molecules associated with apoptotic pathways. For instance, (Sabirzhanov et al. 2020) reported that irradiation-mediated suppression of miR-23a in the brain cortex and hippocampus coincided with elevated levels of Bcl2 family proteins while restoration of miR-23a expression attenuated neuronal apoptosis and cell death. As per (Lin et al. 2011), overexpression of miR-23a reduced H₂O₂-induced cell death and apoptosis in retinal pigment epithelial cells by targeting pro-apoptotic Fas. miR-23a overexpression has also been shown to protect mesenchymal stem cells from TNF-α-mediated apoptosis through regulation of caspase-7 (Mao et al. 2014; Nie et al. 2011; Ruan et al. 2012). In our study too, miR-23a overexpression was found to be neuroprotective in neurotoxin-induced (MPP+, 6-OHDA and Rotenone) cellular PD models by regulating pro-apoptotic BH3-only proteins like NOXA and PUMA. miR-23a was found to directly bind to the 3’UTR of NOXA, thereby down-regulating its expression and preventing the induction of neuronal apoptosis. The study by (Roufayel et al. 2014) further showed that mutation of the miR-23a-binding site in the 3’UTR abolished the miR-23a-mediated repression of the NOXA reporter. Thus, our findings implicate that the reduction of intracellular miR-23a and its enrichment in astrocyte-derived exosomes could be an adaptive stress response, facilitating redistribution of a potentially neuroprotective miRNA to neighbouring neurons during early cellular stress. In the past, other reports have also suggested the protective function of astrocytes under early stages of neurodegenerative stress like AD, by secreting various neuroprotective biomolecules (Saha et al. 2020; Sarkar et al. 2025).

Finally, to investigate the potential clinical relevance of our findings, we examined exosome-associated miR-23a in plasma samples from patients with PD. In plasma derived exosomes, we found a significant increase in miR-23a expression in PD patients, as compared to age-matched controls. However, one caveat is that confirming the cellular (astrocyte) or tissue (brain) origin of these exosomal miR-23a was beyond the scope of our study. Nonetheless, given that the PD patients in our cohort were clinically confirmed to be free of other major co-morbidities, it is plausible that a substantial proportion of circulating miR-23a originates from the brain, and/or astrocytes, as a consequence of progressive neurodegenerative stress. Therefore, our observations should be interpreted as clinical evidence of an association between PD and detectable increase in exosome-derived miR-23a levels in circulating blood.

Overall, our study provides the first evidence that upregulation of CN protein expression in the STR accompanies both astrogliosis and DA neurodegeneration in mouse model of PD. Most importantly, we identified an association between astrocytic Ca²⁺/CN activation and neurotoxin-induced exosomal trafficking of miR-23a, and demonstrated the neuroprotective function of miR-23a by directly regulating the pro-apoptotic BH3-only molecule, NOXA. Finally, increased exosomal miR-23a in plasma from PD patients provides clinical evidence that this pathway has potential translational relevance. Our findings support a model in which astrocytes may respond to early neurotoxic stress by redistributing a potentially neuroprotective miRNA through exosomal communication with neurons.

## 5. Significance and future prospects

Our study clearly indicates that astrocytes respond to PD-associated neurotoxic stress by altering exosome-mediated trafficking of an astrocyte-enriched miRNA, miR-23a, which can protect neurons from apoptotic stress. Moreover, our identification of exosomal miR-23a as a functional, neuroprotective cargo, in the early stages of cellular stress—raises the possibility that exosome-associated miR-23a could serve as an early molecular indicator of PD pathogenesis. Moreover, given that miR-23a overexpression conferred direct neuroprotection against multiple PD-relevant neurotoxins in our study, engineered or endogenously enhanced exosomal delivery of miR-23a could be a potential therapeutic strategy against PD.

More generally, exosome content reflects the molecular status of the originating cell; therefore, from a translational perspective, alterations in exosome cargo profiles, in this case miR-23a, may serve as early indicators of neural dysfunction and ensuing neurodegeneration associated with PD. As such, exosome-based liquid biopsy approaches offer a minimally invasive strategy for diagnosing and monitoring the progression of neurodegenerative disorders such as PD. The ability of certain exosome populations to traverse the blood-brain barrier (BBB) and circulate in body fluids makes them particularly advantageous for biomarker studies as well as broader clinical applications, including nanomedicine (Fais et al. 2016). Exosomes are biocompatible and exhibit low immunogenicity, and when patient-derived, are unlikely to elicit innate or adaptive immune responses (El Andaloussi, Lakhal, et al. 2013; El Andaloussi, Mäger, et al. 2013; Van Niel et al. 2018). Furthermore, owing to their intrinsic capacity for selective cargo loading and delivery, exosomes hold considerable promise as engineered therapeutic carriers for targeted drug delivery to the brain (Bashyal et al. 2022; Rufino-Ramos et al. 2017). For example, NCT03384433 was registered as a Phase I/II clinical trial evaluating allogeneic mesenchymal stromal cell-derived exosomes enriched with miR-124 for acute ischemic stroke, illustrating the emerging clinical translation of miRNA-loaded EV approaches (Van Niel et al. 2022). Building on these precedents, further validation of our findings may position exosome-encapsulated miR-23a as a translational bridge from bench to bedside for both PD biomarker development and therapeutic intervention.

## Funding

This work was supported by institutional funding from the CSIR-Indian Institute of Chemical Biology, Kolkata (CSIR-IICB; P07), Council of Scientific and Industrial Research (CSIR), Ministry of Science and Technology, Govt. of India. PB was supported by the CSIR-Shyama Prasad Mukherjee Fellowship during the course of this project.

## Author Contributions (CRediT format)

PB: Conceptualization, Methodology (cellular, molecular, animal), Formal analysis, Investigation, Data Curation, Writing – Original Draft. KG: Validation (Figure 1), Formal analysis, Writing – Review & Editing. AB: Data curation (clinical). SB: Supervision, Project administration, Writing – Review & Editing, Funding acquisition, Resources.

## Declaration of Competing Interest

The authors declare that they have no known competing interests.

## Data Availability Statement

The data supporting the findings of this study are available within the article and its Supplementary Material. Further inquiries can be directed to the corresponding author upon reasonable request.

## Acknowledgments

We acknowledge the support and facilities provided by the Central Instrument Facility (CIF) and Central Animal facility, CSIR-IICB. The rat PC12 cell line was a kind gift from Prof. Lloyd A. Greene (Columbia University, USA) and the human 1321N1 cell line was kindly gifted by Dr. Sharmistha Banerjee (University of Hyderabad, India). We thank Dr. Rebecca Banerjee (Institute of Neurosciences, Kolkata, India) for assistance in establishing the MPTP mouse model, Dr. Joy Chakraborty (CSIR-IICB) for providing additional mouse tissues and Dr. Suvendra Nath Bhattacharyya’s laboratory (CSIR-IICB) for guidance on miRNA/exosome isolation protocols and access to the NanoSight NS300.

## Declaration of Generative AI and AI-Assisted Technologies in the Writing Process

During the preparation of this work, the authors used ChatGPT (OpenAI) to improve the language and readability of the manuscript. After using this tool, the authors reviewed and edited the content as needed and take full responsibility for the content of the published article.

## Notes

### Competing Interest Statement

The authors have declared no competing interest.

