## Supplemental Figure and Tables for "Exosomal miR-23a release coincides with astrocytic Calcineurin activation, protects neurons from apoptosis by regulating NOXA in Parkinson’s disease models and is enriched in the plasma of patients with Parkinson’s disease"

<u>**Supplementary figures and tables**</u>

**Supplementary Figure:**

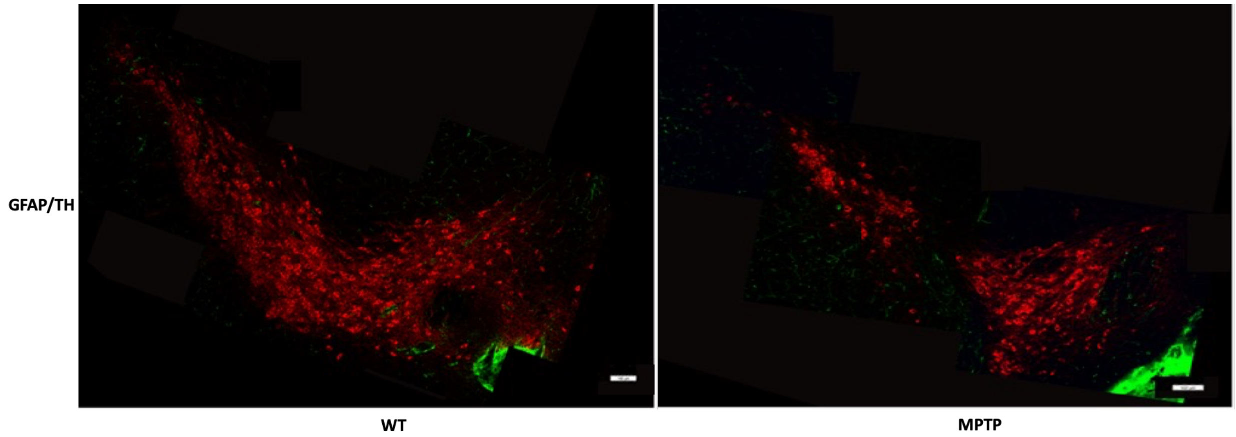


**Figure S1: Decrease in TH positive DA neurons at the SNpc region of MPTP treated mice**. Immuno-histochemical staining of brain sections from SNpc of WT and MPTP-treated mice, imaged under confocal microscope (Leica Sp8 STED microscope). GFAP (green) indicates astrocytes and TH (red) indicate DA neurons. Overlapping images captured at 20X magnification were digitally stitched together using Adobe Photoshop v26.

**Table ST1: List of reagents for miRNA assays**

| **miRNA assay** | **Catalog number (Ambion)** |
|---|---|
| U6 | 4427975 |
| hsa-miR-23a | 4427975 |
| Negative control mimic | 4464058 |
| miR-23a mimic | MC10644 |

**Table ST2: List of primers**

| **mRNA primers** | **Sequence** |
|---|---|
| PUMA forward primer | 5'ACGACCTCAACGCACAGTACGA3' |

| | |
|---|---|
| PUMA reverse primer | 5'CCTAATTGGGCTCCATCTCGGG3' |
| NOXA forward primer | 5'CTGGAAGTCGAGTGTGCTACTC3' |
| NOXA reverse primer | 5'TGAAGGAGTCCCCTCATGCAAG3' |
| GAPDH forward primer | 5'TCAACAGCAACTCCCACTCTT3' |
| GAPDH reverse primer | 5'ACCCTGTTGCTGTAGCCGTAT3' |
| NOXA forward primer (3'UTR cloning) | 5'GACTAGCTCGAGTGACTGCATCAAAAACTTGCATGAGG3' |
| NOXA reverse primer (3'UTR cloning) | 5'CACAGTGCGGCCGCAATTAAAGTGTAAGTCCCTTGAGAG3' |

**Table ST3: List of antibodies**

| Primary antibodies | Catalog number |
|---|---|
| Calcineurin A | Cell Signaling Technology (2614);<br>Novus Biologicals (NBP1-88218) |
| β-Actin | Sigma-Aldrich (A3854) |
| mouse anti-GFAP | Novus Biologicals (NBP1-05197) |
| chicken anti-TH | Abcam (ab76422) |

**Table ST4: Details of human PD patients studied**

| Parameter | Value |
|---|---|
| Number of age-matched controls | 20 |
| Number of PD patients | 25 |
| Number of female PD patients | 07 |
| Number of male PD patients | 18 |
| Average age group of PD patients | 40-70 |
| Number of sporadic PD | 17 |
| Number of familial PD | 08 |
